# Morphine causes distinct changes in the lipidome throughout the brain and body after acute or chronic administration: implications for novel endogenous lipid signaling systems involved in opioid reward and withdrawal

**DOI:** 10.64898/2026.09.17.752401

**Authors:** Taylor J Woodward, Elyse Chafee, Evan Cannon, Clarissa Schmitt, Ken Mackie, Heather B Bradshaw

## Abstract

A growing body of evidence demonstrates that signaling pathways of endogenous lipids (endolipids) modulate the reinforcing properties of opioids including reward and withdrawal. Many of these studies are focused on the endocannabinoid (eCB) system, and its primary eCB ligands, Anandamide (AEA) and 2-arachidonoylglycerol (2-AG). The central hypothesis is that modulation of the eCB system through eCB receptors and enzymes may improve therapeutic outcomes for opioid use disorder (OUD). However, outcomes in preclinical and clinical studies using the low efficacy, CB1/CB2 orthosteric agonist, THC, found limited to no effectiveness, suggesting that targeting this aspect of CB1/CB2 is not a useful therapeutic tool for OUD. Previous studies finding that genetic deletion or pharmacological inhibition of endolipid-regulating enzymes (including eCBs) can alter behavioral sensitivity to opioids provide insight into an alternative approach. If opioid use dysregulates a wide range of endolipid biosynthesis and metabolism, then a clearer understanding of these changes, especially in signaling ligands, will provide a novel avenue to both understand the underlying physiological changes with opioid use as well as providing novel targets for therapeutic interventions. In this study, we tested the hypothesis that if morphine dysregulates multiple endolipid signaling systems in the brain and body, and this dysregulation evolves over time, then these differential effects will be measurable by changes in endolipid levels. Using an Acute (30 minutes post injection 20mg/kg) and a Chronic paradigm (5 days, twice daily, 20mg to 100mg/kg escalating dose) we measured 100 targeted endolipids in 8 brain regions, plasma, liver, and feces in male mice. In the Acute condition, we found that the most screened endolipids were changed in the striatum (42%), while the fewest were changed in the thalamus (19%) and 38% in the plasma. In the Chronic condition, 79% of plasma endolipids were changed. The highest level of change in the CNS was in the cortex (39%). Levels of AEA and 2-AG were largely unchanged; however, levels of the *N*-acyl GABAs, *N*-acyl valines, *N*-acyl taurines, and specific bile acids (*e.g.* DCA, TCA) showed the most dynamic changes by treatment group. These results provide information on novel endolipid signaling pathways that may contribute to the unwanted side effects of opioids, such as dependence and withdrawal, and provide novel avenues for the development of therapeutic strategies.

## Introduction

Developing effective therapies for opioid use disorder (OUD) is challenging due to the complexity of the broad range of physiological adaptations that occur in the brain and body during chronic exposure to opioids. While medications such as buprenorphine, methadone, and naloxone have emerged as useful strategies to treat components of OUD, including withdrawal, craving and overdose, they face challenges in terms of acceptance, accessibility, cost, and abuse liability [1]. An understanding of the wider range of physiological systems that are affected by opioid use, including endogenous lipid signaling, will serve in the development of novel therapeutic strategies and allow for more targeted approaches.

A growing body of evidence shows that signaling pathways of endogenous lipids (endolipids) modulate the reinforcing properties of opioids including tolerance, reward, and withdrawal [2-9]. Many of these studies focus on the endocannabinoid (eCB) system, and its primary eCB ligands, Anandamide (AEA) and 2-arachidonoylglycerol (2-AG), with the hypothesis that eCB system signaling through the CB1 and CB2 receptors are the targets involved in therapeutic outcomes. However, outcomes in preclinical and clinical studies using the CB1/CB2 orthosteric agonist, Δ9-Tetrahydrocannabinol (THC), showed mixed effects on OUD outcomes and limited effectiveness overall [10-14]. This suggests that targeting CB1/CB2 receptors with orthosteric agonists is not sufficient as a standalone therapeutic for OUD. An alternative strategy for treating OUD that has been explored in preclinical studies is elevating levels of AEA or 2-AG by inhibiting their metabolic enzymes, Fatty Acid Amide Hydrolase (FAAH) and Monoacylglycerol Lipase (MAGL) [3, 5, 7]. This strategy, too, has inconsistent effectiveness.

One explanation for the lack of overall effectiveness of targeting CB1 and eCB metabolic enzymes is that AEA and 2-AG are only 2 members of a much larger group of hundreds of endolipid congeners with a wide range of signaling pathways [15-18][15, 18, 19]. These endolipids, as a family of signaling molecules, serve as key regulators of diverse physiological processes throughout the brain and body including neurotransmission, pain perception, sleep, immune response and thermoregulation via actions at various receptors, including G-protein coupled receptors (GPCRs), ion channels, and transcription factors [15, 16]. Therefore, if a wider range of endolipid biosynthesis and metabolism is dysregulated with opioid use, then the signaling system outcomes are potentially exponential and bringing those signaling systems back into homeostasis will require more than the activity at single receptors.

To test the hypothesis that opioids broadly dysregulate endolipids, we chose a targeted lipidomic approach to focus on specific endolipid signaling systems related to eCBs (100 endolipids in total screened) [15]. To capture temporal components, we focused on 2 time points: 1) Acute, which is single dose of morphine (20 mg/kg, i.p.) with tissue collected 30 minutes post injection, and 2) Chronic, which collects tissue 24 hours after a 5-day, twice-daily, escalating dosing paradigm. This paradigm is well characterized to produce behavioral signs of spontaneous opioid withdrawal [20-22]. We tested the hypothesis that if morphine causes dysregulation of multiple endolipid signaling systems in the brain and body, and this changes over time, then these differential effects will be measurable by changes in specific endolipid levels. This information will guide us to a better understanding of how endolipid signaling changes across the brain and body during opioid use and withdrawal, which will allow us to target specific endolipid signaling and metabolic pathways beyond the canonical eCB signaling systems to identify novel therapeutic targets.

## Methods

### Animals

All procedures were approved by the Institutional Animal Care and Use Committee at Indiana University Bloomington. 32 male C57BL6/J mice (11 weeks old) were purchased from Jackson Laboratories (Bar Harbor, ME) and allowed to acclimate to housing facilities for 1 to 2 weeks prior to experimental treatments. Mice were housed 2 per cage under a standard 12-hour light/dark cycle (lights on at 8:00 AM) with *ad libitum* access to food and water.

### Drug Administration

Morphine sulfate (10 mg/mL), purchased from Henry Schein (Mellvile, NY), was diluted in sterile saline to appropriate concentrations for injections. Mice were randomly assorted to receive saline or morphine by cages, and morphine was administered intraperitoneally at a volume of 10 µL/g body weight. One cohort of mice (Acute) received a single injection of morphine (20 mg/kg) or saline 30 minutes prior to sacrifice/tissue collection (n=8 saline and n=8 morphine). A second cohort of mice (Chronic) received twice daily escalating doses of morphine (or equivalent volume saline) over the course of 5 days in the morning (∼8:30 AM) and afternoon (∼4:00 PM) under the following dosing regimen: Day 1-20 mg/kg, Day 2-40 mg/kg, Day 3-60 mg/kg, Day 4-80 mg/kg, and Day 5-100 mg/kg. Mice treated with chronic morphine were sacrificed 24 hours after the final injection of morphine (100 mg/kg) on Day 6, a time point that has widely been documented with this dosing regimen to produce behavioral signs of spontaneous opioid withdrawal and lasting alterations in neurophysiology [20-22].

### Tissue collection

After acute or chronic treatment with morphine or saline, mice were rapidly decapitated without anesthesia. Brains were rapidly removed, placed in a 15 mL conical tube, and flash frozen in liquid nitrogen. The right medial lobe of the liver was dissected, placed into a tube, and flash frozen in liquid nitrogen. Fecal samples were collected either from expulsion during sacrifice, or from the colon. In cases where feces were not expelled during sacrifice, the most distal fecal bolus was removed from the colon. Fecal samples were placed in a microcentrifuge tube and frozen in liquid nitrogen. Trunk blood was collected after decapitation in heparinized tubes and placed on ice until centrifugation at 1950 x g for 12.5 minutes at 4°C. Plasma aliquots and all other tissue samples were stored in a -80°C freezer until further processing.

Brains were dissected into gross anatomical regions as described previously [18, 23]. To summarize, brains were briefly thawed on a glass plate (∼5 minutes) on ice. The hypothalamus (HYP), striatum (STR), hippocampus (HIPP), cortex (CTX), cerebellum (CER), midbrain (MID), thalamus (THAL), and brainstem (STEM) were quickly separated into tubes that were flash frozen in liquid nitrogen. A small piece of the liver (∼25mg) was also dissected in this manner on ice prior to tissue processing.

### Lipid extraction and partial purification

Lipids were extracted and partially purified as previously performed [24, 25]. In brief, tissue samples (brain parts, liver, feces, or 50 uL of plasma) were placed into 2 mL of HPLC grade methanol (MeOH) on ice and spiked with 10 µL of a mix of deuterium labeled lipids purchased from Cayman Chemical (Ann Arbor, MI, USA) including *N*-arachidonoyl ethanolamine-d8 (1 µM), *N*-palmitoyl ethanolamine-d5 (100 nM), *N*-oleoyl ethanolamine-d4 (100 nM), *N*-arachidonoyl serine-d8 (1 µM), d4-arachidonoyl taurine (1 µM), prostaglandin E2-d4(1 µM), cortisol-d4 (1 µM), deoxycholic acid-d4 (1 µM), and morphine-d3 (1 µM). Solid tissues (brain, liver, feces) were incubated on ice in the dark for 2 hours, sonicated (∼20 seconds) and centrifuged at 19,000g for 20 minutes at 20°C. Plasma samples were incubated in the dark on ice for 30 minutes prior to centrifugation under the same conditions. After centrifugation, the 2 mL of methanolic supernatant was added to 8 mL of HPLC water to create a 20% organic solution. This was passed dropwise through Bond Elut solid phase extraction (SPE) columns (Agilent, Santa Clara, CA) previously activated with 5 mL of HPLC grade methanol and 2.5 mL of HPLC grade water. Columns were washed with 2.5 mL water followed by 1.5 mL of 40% methanol. 1.5 mL elutions of 75% followed by 100% methanol were stored at -80°C.

### High-Performance Liquid Chromatography and Mass Spectrometry

A library of 100 endolipids and morphine was screened from each sample via a SCIEX 7500 mass spectrometer (SCIEX, Marlborough, MA, USA) coupled to a Shimadzu LC system LC-40DX3 (Shimadzu, Kyoto, Japan). Mobile phase A was 1 mM ammonium acetate in 20% LCMS grade MeOH/80% LCMS grade water, and mobile phase B was 1 mM ammonium acetate in 100% LCMS-grade MeOH. Mobile phase flow rate was 0.2 mL/minute. Lipids were detected from a 10 µL injection of each sample’s partially purified extract (75% and 100% elution) using an electrospray ionization (ESI) combined positive/negative mode multiple reaction monitoring (MRM) method previously optimized for each analyte. A full list of endogenous lipids and deuterium-labeled internal standard, along with parent/fragment mass for each is presented in Supplemental Figure 1. Sample concentrations of each analyte were quantified by fitting each sample’s peak area to a calibration curve of diluted standard using SCIEX OS version 3.4 (SCIEX, Marlborough, MA, USA) and were adjust for recovery and normalized to each tissue’s mass or volume (for plasma) to create a final moles/g or moles/mL concentration for each sample.

### Statistical Analysis

Morphine concentrations were analyzed using one-way ANOVA, and weight loss during chronic morphine dosing was analyzed using a two-way ANOVA. Lipidomic data were analyzed using a two-tailed t-test for each tissue using a custom script written in Python (version 3.12.4), which calculated a p value, fold change, and Cohen’s D for each comparison. A summary heatmap was also generated, with statistically significant (p<.05) denoted with darker colors and trending comparisons (p value between 0.1 and 0.05) with lighter colors.

Arrows depicting directionality of change (increase or decrease) of morphine compared to saline are also included in this summary heatmap, where one arrow corresponds to a 1-1.49 fold change, 2 arrows corresponds to a 1.5-1.99 fold change, 3 arrows corresponds to a 2-2.99 fold change, 4 arrows corresponds to a 3 to 9.99 fold change, and 5 arrows corresponds to a 10+ fold change as described previously [24, 25]. Treatment comparisons with fewer than 3 detected peaks in a group are labeled as “Below Analytical Level (BAL), while Below Detection Level (BDL) is used when no peaks were detected in any group for a comparison. Figures were created with GraphPad Prism (GraphPad, San Diego, CA, USA).

## Results

### Morphine was detected in the CNS after Acute and Chronic administration

The experimental design schematic is shown in Fig 1A to visualize injection timing, tissue collection, and sample processing/analysis workflow. The first level of analysis was to determine the level of morphine in each tissue type and to validate that morphine was present throughout the brain and body. In the Acute condition (Fig 1B), 30 minutes after the 20 mg/kg i.p. injection of morphine, morphine was present in all tissues screened. In the CNS, levels were ∼2-fold higher in the midbrain (MID) compared to all other CNS brain regions, which had equivalent levels. In the periphery, morphine levels in the liver were equivalent to the CNS levels (apart from MID), plasma levels were ∼3-fold higher, and levels in the feces were ∼10-fold higher. Morphine was still detectable in all tissues 24 hours after the final 100 mg/kg injection of morphine in the Chronic condition (Fig 1C), with levels in the CNS and liver being equivalent and ∼2-fold higher than levels in plasma. The level of morphine in the feces in the Chronic condition was ∼1000-fold higher than all other tissues screened.

**Figure 1:**
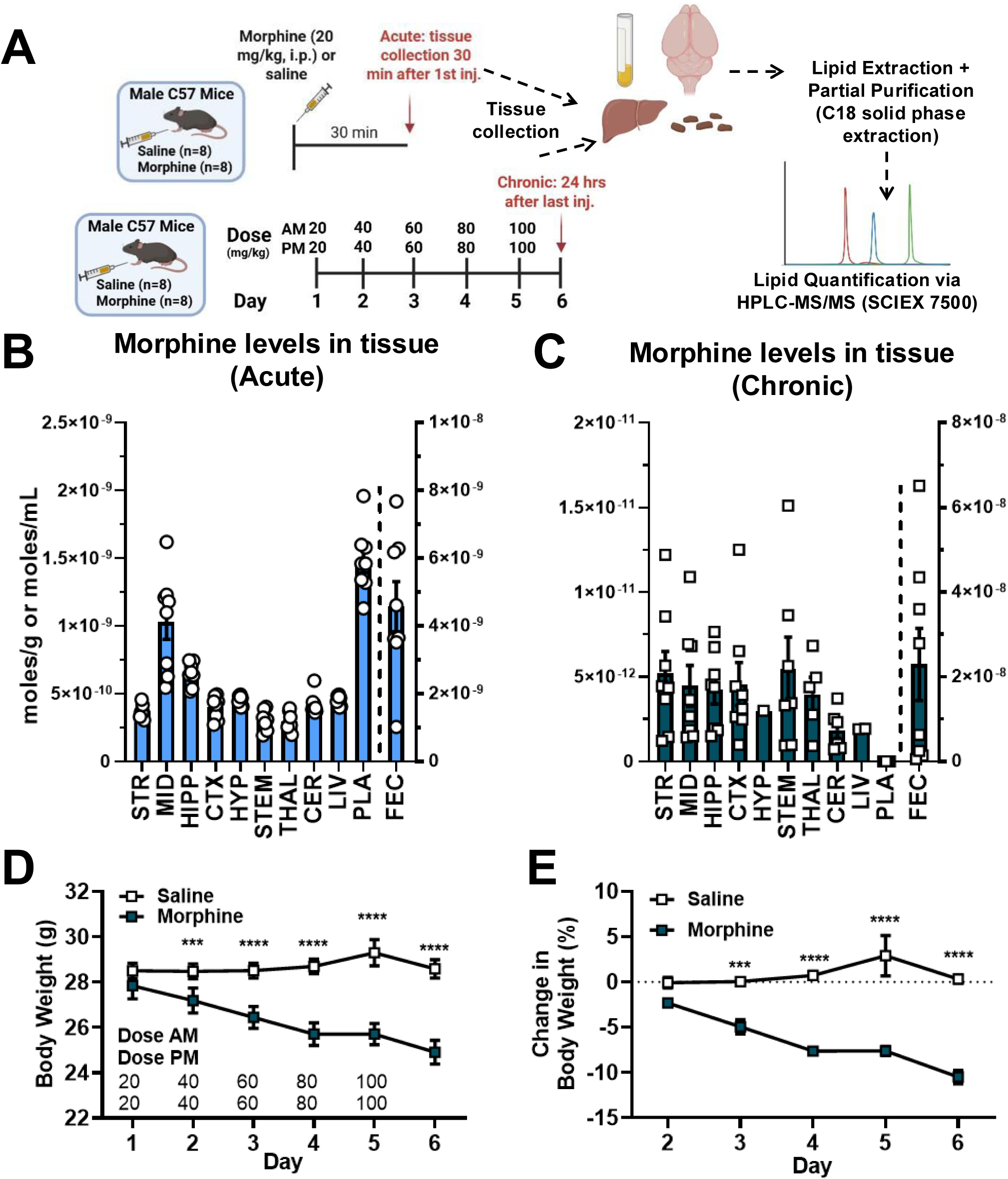
Acute and Chronic morphine treatment paradigm shows levels of systemic morphine throughout the brain and body and changes in body weight. A) Experimental schematic showing injection schedule and timepoint of sacrifice for mice in acute and chronic conditions. B) Concentrations of morphine in CNS and peripheral tissues after acute injection of morphine (20 mg/kg, i.p.) [One way ANOVA: F(10,67)=29.62, P<.0001]. C) Concentrations of morphine 24 hours after the final injection of chronic escalating morphine in CNS and peripheral tissues [One way ANOVA: F(10,58)=5.308, P<.0001]. Fecal levels of morphine plotted on a separate y axis to display tissues on the same graphs. D) Raw body weight (in grams) of mice receiving chronic saline or morphine during chronic injections E) Change in body weight (normalized to pre-injection day 1) of mice receiving chronic saline or morphine. [Two Way ANOVA for D: Time F(5,35) = 11.52, P<.0001; Drug F(1,7)=11.92, P=.0107; Time x Drug F(5,35)=19.42, P<.0001][ Two Way ANOVA for E: Time F(4,28) = 6.660, P=.0007; Drug F(1,7)=75.69, P<.0001; Time x Drug F(4,28)=12.20, P<.0001] All data are presented as mean +/-SEM. Bonferroni’s post hoc comparisons of morphine compared to saline: *** p<.001, **** p<.0001. Note that in some instances error bars are too small to visualize due to low variability. Tissue Abbreviations: STR=striatum, MID=midbrain, CTX= cortex, HYP= hypothalamus, STEM=brainstem, THAL=thalamus, CER= cerebellum, PLA=plasma, LIV=liver, FEC=feces

Representative chromatograms of the morphine standard, along with examples of morphine detected in the CNS and plasma of mice treated with chronic morphine are provided in Supplemental Figure 2. Chronic morphine treatment significantly decreased body weight, presented in grams (Fig 1D) as well as percent change from baseline (Fig 1E).

### Morphine significantly changes levels of a wide range of lipids in both the Acute and Chronic conditions

Three key levels of analysis (p-value significance, fold-change, and Cohen’s D-value effect size) showed that morphine treatment induced significant tissue-specific changes in endolipid levels after Acute and Chronic morphine treatment *(See Methods for details; See Supplemental Figure 3 for comparisons of each of these levels of analysis*). Supplemental Figures 3C and 3D show that the Log2 Fold-change values are positively correlated (p<0.0001) with Cohen’s D values for Acute (r=.79) and Chronic (r=.85) comparisons. Because the fold-change values and the Cohen’s D effect values are highly correlated and because the fold-change values provide a higher degree of granularity and the ability to plot against p values, we use fold-change values to denote magnitude of effect here. Volcano plot visualizations summarize the effects of Acute and Chronic morphine (Fig 2A-B) with peripheral tissues plotted in grey, and CNS tissue plotted in pink. For each comparison (vehicle compared to morphine treatment), log2 fold-change is plotted on the x-axis and log10 p-value plotted on the y-axis. This allows for identification of comparisons using the magnitude of change (fold-change, Cohen’s D), statistical significance (p-values), and directionality of effect (positive values indicating increases and negative values are decreases) for specific endolipid species across tissue types.

**Figure 2:**
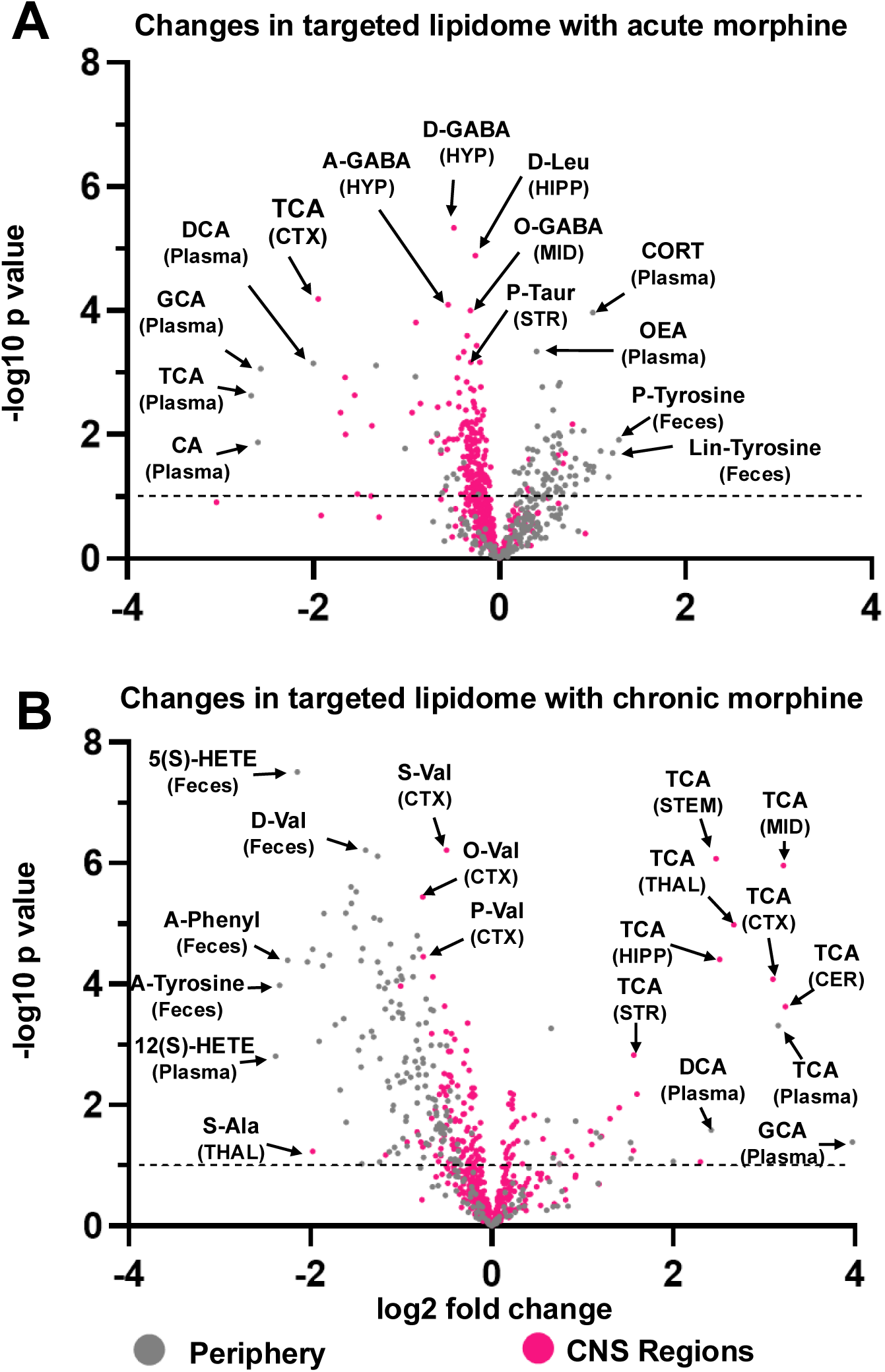
Volcano plot summaries of changes in endolipid levels after morphine treatment. Volcano plots visualize fold-change and p-value of each endolipid screened in each tissue from (A) acute morphine treatment (20 mg/kg; tissue collection at 30 minutes) compared to acute saline, (B) chronic morphine (5-days increasing dose with final dose 100mg/kg with tissue collection at 24 hours post final dose) compared to chronic saline. The x-axis displays log2 fold change in endolipid concentration as a function of endolipid levels measured in the morphine treatment compared to those in the saline treatment (*i.e.* 1 denotes that the specific morphine treatment doubled an endolipids concentration, and -1 denotes that the specific morphine treatment halved an endolipid’s concentration). The y-axis displays -log10 p values for the comparisons of levels in the morphine treatment compared to saline (*i.e.* values above 1 on the y-axis represent a p value of p<.1, whereas those with p <0.05 begin to be represented at ∼1.3 on the scale). On each plot, comparisons performed in CNS regions are represented in pink, and comparisons for peripheral tissues (plasma, liver, and feces) are represented in grey. Endolipid abbreviations: CA=cholic acid, GCA=glycocholic acid, TCA=taurocholic acid, DCA=deoxycholic acid, A-GABA=*N-*arachidonoyl gamma-aminobutyric acid, D-GABA=*N*-docosahexaenoyl gamma-aminobutyric acid, O-GABA=*N-*oleoyl gamma-aminobutyric acid, D-Leu=*N*-docosahexaneoyl leucine, P-taur=*N*-palmitoyl taurine, CORT=corticosterone, OEA=*N*-oleoyl ethanolamine, P-tyrosine=*N*-palmitoyl tyrosine, Lin-tyrosine=*N*-linoleoyl tyrosine, S-ala=*N*-stearoyl Alanine, 12(S)HETE=12(S)-Hydroxyeicosatetraenoic Acid, A-Tyrosine= *N*-arachidonoyl tyrosine, A-Phenyl=*N*-arachidonoyl phenylalanine, 5(S)HETE=5(S)-Hydroxyeicosatetraenoic Acid, D-Val=*N*-docosahexaenoyl valine, S-Val=*N*-stearoyl valine, O-Val=*N*-oleoyl valine, P-Val=*N*-palmitoyl valine Tissue Abbreviations: STR=striatum, MID=midbrain, CTX= cortex, HYP= hypothalamus, STEM=brainstem, THAL=thalamus, CER= cerebellum

The overall effect on total significant changes across all tissues for this targeted lipidome changed as a function of treatment and body region (Supplemental Figure 4-7). After Acute morphine (Fig 2A), there is a clear divergence of the directionality of change of endolipids with significant changes. In the CNS, ∼29% decreased and ∼1% increased, whereas in the periphery only ∼7% decreased but ∼30% increased. Chronic morphine (Fig 2B) caused an almost complete shift to decreases (∼55%) verses increases (∼6%) in the periphery (∼55%) while most changes in the CNS remained decreases (∼17%) verses increases (∼7%).

Figures 3A-B summarize morphine’s effects as the percent of endolipids significantly changed by tissue type. Tissues screened here are arranged in descending order by overall percent change of Acute morphine where the striatum (STR) had the highest percent of endolipids changed (42%) and the thalamus (THAL; 19%) the least. In the Chronic condition, only 15% of striatal endolipids changed, and the tissue with the most significant changes was plasma (79%), followed closely by feces (69%). The CNS tissue in the Chronic condition with the most changes was the cortex (39%).

Figure 3C shows the top 10 endolipids with highest instance of change across tissues for each treatment paradigm. Only one endolipid, deoxycholic acid (DCA; Fig. 3C, 6F-J), was in the top 10 most affected endolipids in both treatment paradigms (Acute 70%; Chronic 63%). In the Acute condition, *N*-acyl gamma-aminobutyric acid (*N*-acyl GABA) endolipids were significantly reduced by morphine across the CNS (Fig. 4A, 4C). *N*-arachidonoyl GABA was the only endolipid that was altered by morphine in every tissue in which it was detected at analytical levels (Fig. 3C), which was only in the CNS as *N*-arachidonoyl GABA was not detected in the peripheral tissues in our screen (Fig 4A). The top spot with significant differences in 10 out of 11 tissues (90%) in the Chronic condition (Fig. 3C), Taurocholic acid (TCA; Fig. 6A-6E), a taurine-conjugated bile acid, was present in all tissue measured here.

**Figure 3:**
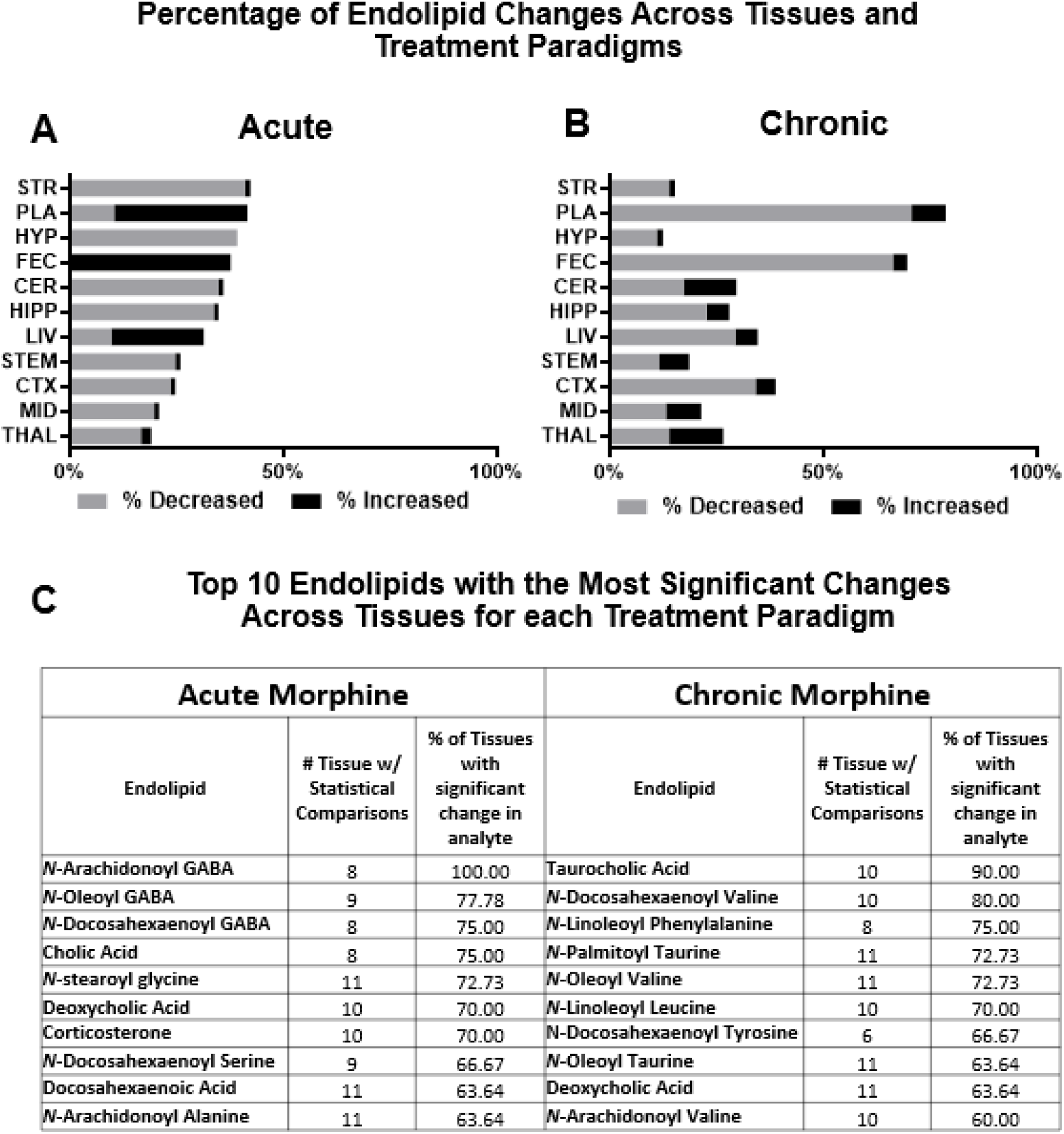
Direction of change in Endolipid levels after morphine treatment varied by tissue type and treatment condition. Percentages of endolipids altered by acute (A) and chronic (B) morphine across tissue types. For each tissue, percentage of significant decreases in endolipid levels is denoted with a pink bar, and percentage of significant increases is denoted in dark blue. For both conditions, tissues are arranged in descending order by overall percent change of acute morphine. C) A table summarizing the top 10 endolipids most frequently modulated across tissues by either Acute (left) or Chronic (right) morphine treatment. For each treatment type, the far-left column in the table lists the individual endolipid, the middle column lists the number of tissues in which the endolipid was detected at analytical levels (see Methods), and the far-right column is the percentage endolipids detected that were significantly changed. Tissue Abbreviations: STR=striatum, MID=midbrain, CTX= cortex, HYP= hypothalamus, STEM=brainstem, THAL=thalamus, CER= cerebellum, PLA=plasma, LIV=liver, FEC=feces

**Figure 4:**
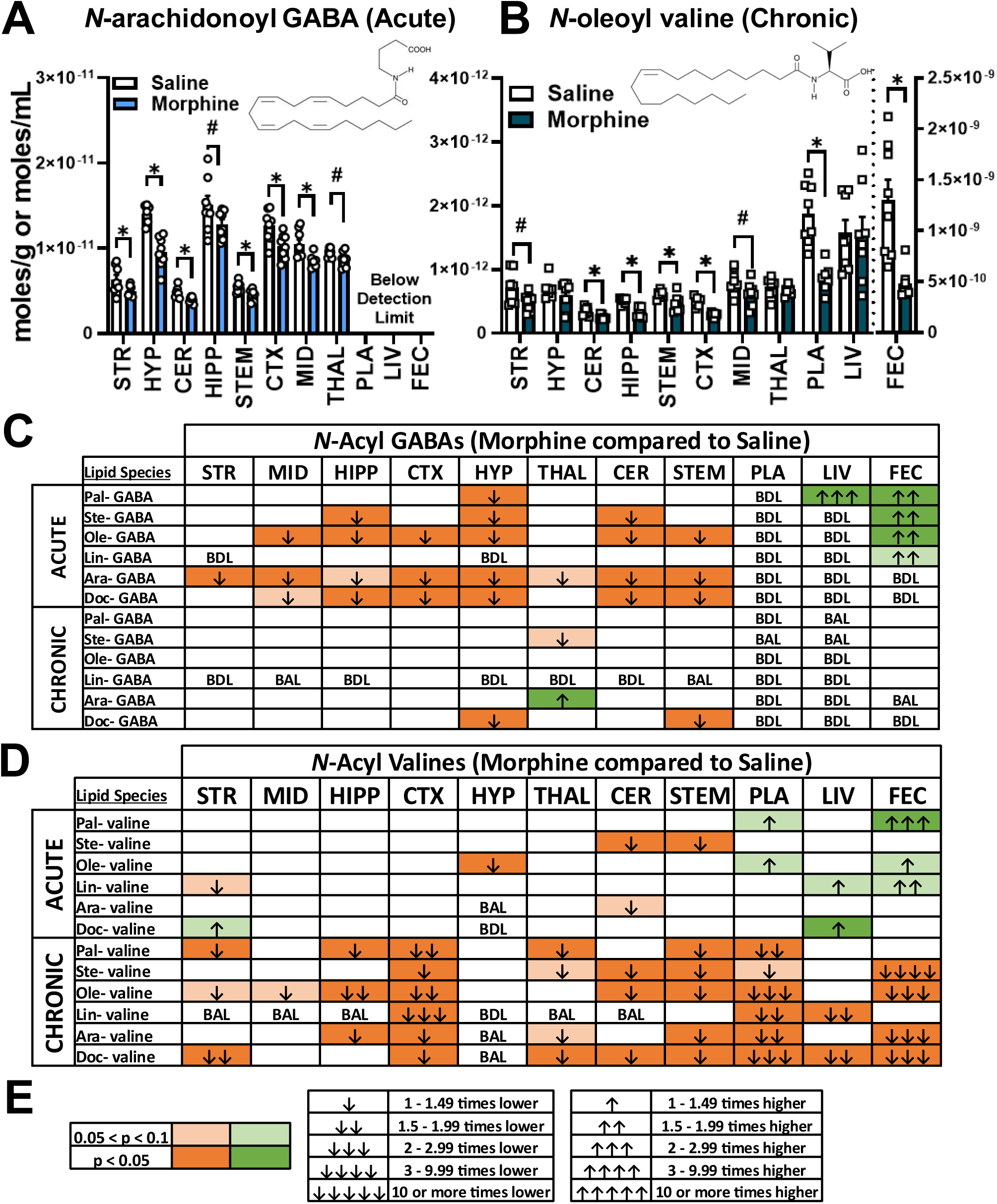
Effect of acute and chronic morphine on levels of *N*-acyl GABA and N-acyl valine endolipid families. A) Concentrations in moles/gram of *N*-Arachidonoyl GABA across tissues after acute treatment (not detected in peripheral tissues). B) Concentrations of *N*-oleoyl valine after chronic morphine treatment in moles/gram for tissues or moles/mL plasma. Fecal concentrations of both representative lipids are plotted on a separate y-axis on the right due to differences in tissue concentrations. Two tailed t-test: #=p value between .05 and .01, *=p<.05. Data are presents as the mean +/-SEM with n=8 per treatment group. C) Heatmap summarizing effects of acute and chronic morphine on *N*-acyl GABAs. D) Heatmap summarizing effects of acute and chronic morphine on *N*-acyl valines. E) Key to heatmap: light shading represents 0.1>p>.05, dark shading represents p<.05 for a given comparison. Green boxes with upward arrows depict that morphine increased levels of a lipid within a tissue, whereas orange boxes with down arrows depict the opposite. Acyl Group Abbreviations: Pal=*N-*palmitoyl, Ste=*N*-stearoyl, Ole=*N*-oleoyl, Lin=*N*-linoleoyl, Ara=*N*-arachidonoyl, Doc=*N*-docosahexaenoyl Tissue Abbreviations: STR=striatum, MID=midbrain, CTX= cortex, HYP= hypothalamus, STEM=brainstem, THAL=thalamus, CER= cerebellum, PLA=plasma, LIV=liver, FEC=feces Heatmap key: Heatmaps were generated to summarize the effect of morphine on each lipid species in each tissue. Light green or light orange boxes depict a P value between 0.1 and 0.05, while dark green or dark orange depict a P value below 0.05. Arrows summarizing fold change of morphine compared to saline are presented as an increase(↑) or decrease(↓). Number of arrows correspond with magnitude of fold change. One arrow: 1-1.49 fold difference; two arrows: 1.5-1.99 fold difference; three arrows 2-2.99 fold difference; four arrows 3-9.99 fold difference, five arrows 10+ fold difference.

Levels of 2 additional taurine-related endolipids, *N*-palmitoyl taurine (Fig. 5A-5E) and *N*-oleoyl taurine (Fig. 5F) were significantly changed in 72% and 63% of tissues respectively. Of note, the levels of the eCBs (Supplemental Fig. 8), AEA and 2-AG were largely unchanged. In the Acute condition (Supp. Fig. 8A-B), AEA decreased in the hypothalamus and 2-AG had no changes across all tissues. In the Chronic condition (Supp. Fig. 8C-D), AEA and 2-AG were both significantly decreased in plasma. 2-AG was also decreased in the hippocampus and the feces. Because other endolipids were more significantly modified in both the Acute and Chronic conditions here, our results and discussion will focus on those endolipid species and not on these canonical eCB ligands.

**Figure 5:**
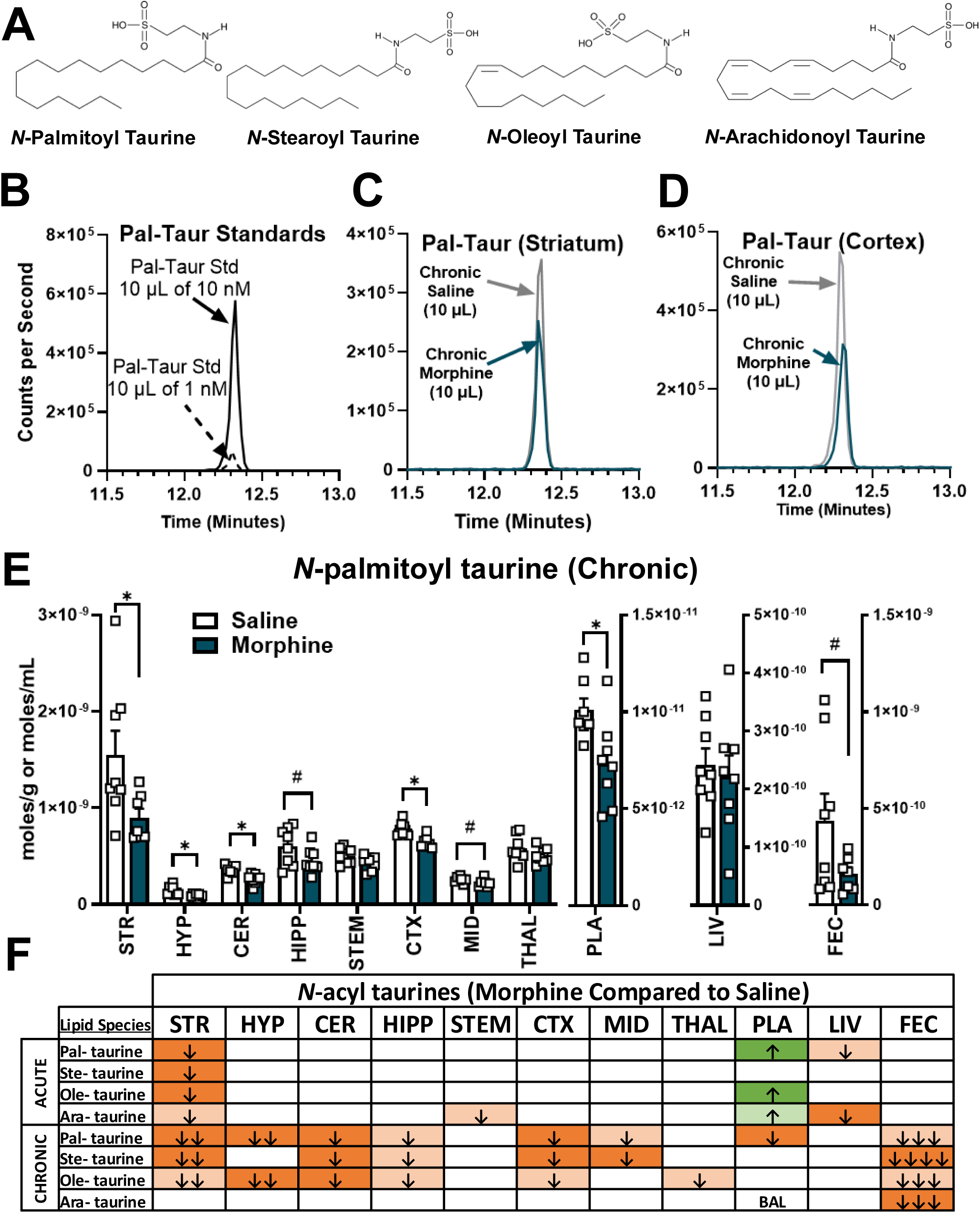
Effects of morphine on levels of *N*-acyl taurines in the CNS and periphery. Chemical structures are displayed for *N*-acyl taurines (NATs) screened in the present study B) Representative chromatogram of two different concentrations of *N*-palmitoyl taurine standards (10 µL injection of 1 and 10 nanomolar). C) Representative chromatograms of *N*-palmitoyl taurine quantified from the striatum (STR) of a mouse treated with Chronic saline (grey; 10 µL injection) or Chronic morphine (dark blue; 10 µL injection). D) Representative chromatograms of *N*-palmitoyl taurine quantified from the cortex (CTX) of a mouse treated with chronic saline (grey; 10 µL injection) or chronic morphine (dark blue; 10 µL injection). E) Bar graph visualizing effect of morphine on levels of *N*-palmitoyl taurine across tissues in moles/g for tissues and moles/mL for plasma F) Heatmap summarizing effect of acute and chronic morphine on levels of NATs in CNS and peripheral tissues. Two tailed t-test: #=p value between .05 and .01, *=p<.05. Data are presents as the mean +/-SEM with n=8 per treatment group. Acyl Group Abbreviations: Pal=*N-*palmitoyl, Ste=*N*-stearoyl, Ole=*N*-oleoyl, Ara=*N*-arachidonoyl Tissue Abbreviations: MID=midbrain, HYP= hypothalamus, STEM=brainstem, THAL=thalamus, CER= cerebellum, PLA=plasma, LIV=liver, FEC=feces

**Figure 6:**
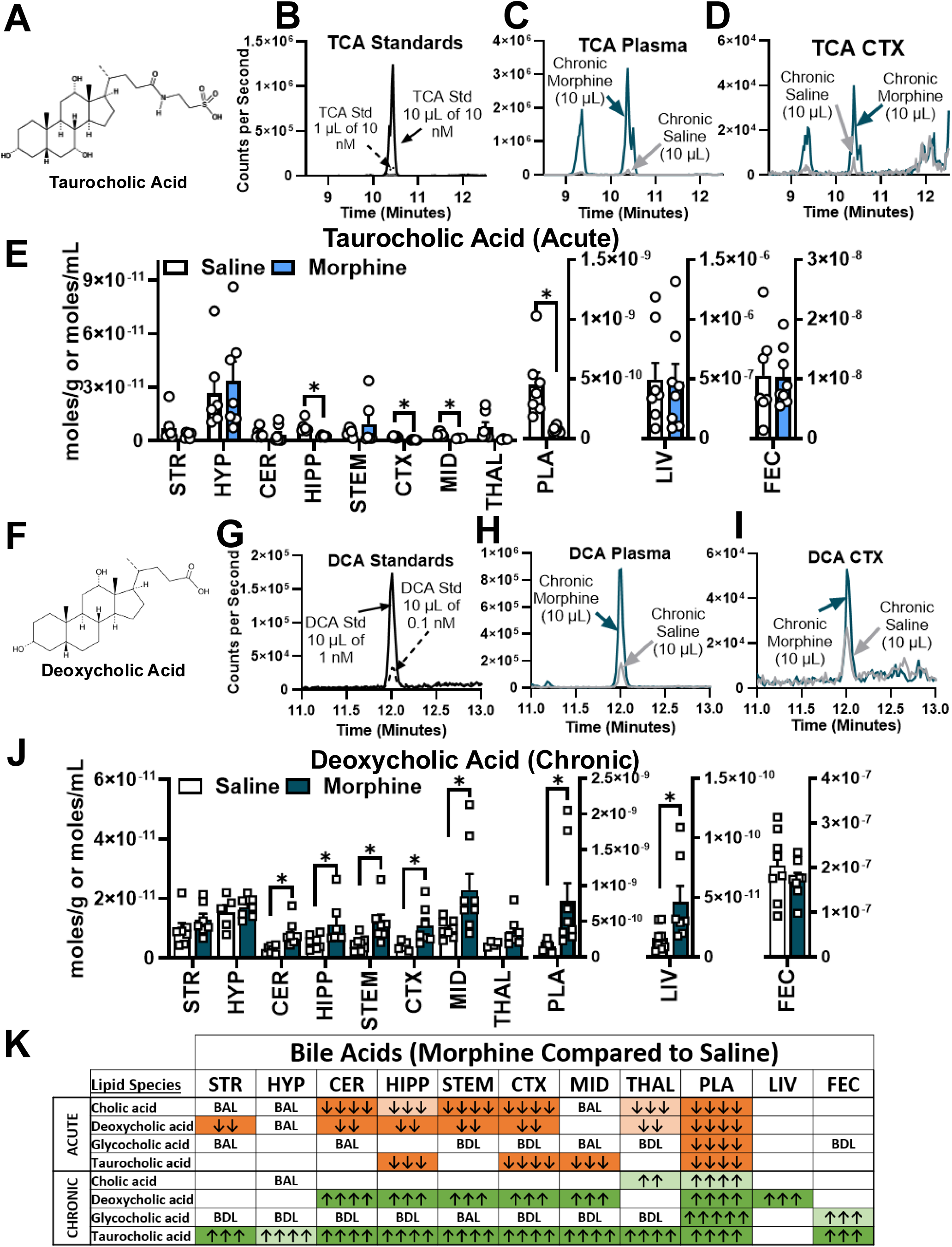
Detection of bile acids in the CNS and effects of acute/chronic morphine on bile acid levels. Chemical structure of taurocholic acid (TCA) B) Chromatogram of 2 different concentrations of a taurocholic acid standard (1 µL and 10 µL of 10 nanomolar). C) Representative chromatograms of TCA in plasma from mice treated with chronic saline (grey) or chronic morphine (dark blue). D) Representative chromatograms of TCA in the cortex (CTX) of mice treated with chronic saline or morphine. E) Concentrations of TCA in tissues from chronic saline or chronic morphine treated mice. F) Chemical structure of deoxycholic acid (DCA). G) Chromatogram of 2 different concentrations of a taurocholic acid standard (10 µL of 0.1 and 1 nanomolar). H) Representative chromatograms of DCA in plasma from mice treated with chronic saline or chronic morphine. D) Representative chromatograms of DCA in the cortex (CTX) of mice treated with chronic saline or morphine. E) Concentrations of DCA in tissues from chronic saline or chronic morphine treated mice. K) Heatmap summarizing effects on bile acids evaluated in the present study. Two tailed t-test: #=p value between .05 and .01, *=p<.05. Data are presents as the mean +/-SEM with n=8 per treatment group. Tissue Abbreviations: STR=striatum, MID=midbrain, HYP= hypothalamus, STEM=brainstem, THAL=thalamus, CER= cerebellum, PLA=plasma, LIV=liver, FEC=feces

### Acute and Chronic Morphine differentially alter levels of *N*-Acyl GABAs and *N*-Acyl Valines

Acute morphine significantly decreased levels of members of the *N*-acyl GABA (NAGABA) family throughout the CNS. As shown in Figure 4A, Acute morphine significantly decreased levels of *N*-arachidonoyl GABA in every CNS tissue but was below detection limits in the periphery. Figure 4C provides a heatmap summary of morphine’s effects on NAGABAs, demonstrating that acute morphine similarly reduced levels of *N*-oleoyl GABA and *N*-docosahexaenoyl GABA. *N*-linoleoyl GABA was the only member of this family unaffected by acute morphine in the CNS. However, *N*-linoleoyl GABA together with *N*-oleoyl GABA, *N*-stearoyl GABA, and *N*-palmitoyl GABA was significantly increased in the feces. By contrast, Chronic morphine had little overall effect on levels of NAGABAs, though one of the few notable changes was that chronic exposure to morphine increased levels of *N*-arachidonoyl GABA in the thalamus.

Chronic morphine decreased levels of *N*-acyl valines across the CNS and periphery. As shown in Figure 4B, levels of *N*-oleoyl valine (a representative of this family) was decreased in 6/8 brain regions, plasma, and feces. Figure 4D shows a comparison heatmap for the *N*-acyl valines (NAVALs), which illustrates the dramatic shift in responses between morphine treatment conditions. Overall, Acute morphine showed few changes in NAVALs in the CNS, however, significant increases in peripheral tissues. By contrast, Chronic morphine caused significant reductions in all NAVALs throughout the CNS and periphery, especially in the cortex where all 6 members of this family were decreased.

### Acute and chronic morphine treatment differentially altered levels of *N*-acyl taurines in the CNS and periphery

Figure 5A displays chemical structures and differences in acyl chain between different *N*-acyl taurines (NATs) evaluated in the present study. Given that few studies to date evaluate the NATs in the CNS, in Figure 5 we show chromatograms of representatives of NATs to illustrate the relative abundance of these endolipids. Figures 5B shows representative chromatograms of an *N*-palmitoyl taurine standard (1 nM and 10 nM). Figures 5C and 5D are representative chromatograms of the striatum (STR) and cortex (CTX), respectively, of mice treated with chronic saline (grey) or chronic morphine (dark blue). As seen in bar graph visualizations in Figure 5E, chronic morphine decreased levels of *N*-palmitoyl taurine in 6 of 8 brain regions, in addition to plasma and feces. A summary heatmap in Figure 5F illustrates the differential effects of Acute and Chronic morphine on NAT levels in the CNS and periphery. Acute morphine only affected NATs in the striatum, causing significant decreases in all species. Likewise, NAT levels significantly decreased in liver. However, NAT levels in plasma were significantly increased. Chronic morphine caused significant reductions in each NAT species across the CNS except, *N*-arachidoyl taurine. Similarly, fecal levels of all four NATs were significantly reduced.

### Acute morphine decreased, and Chronic morphine increased levels of bile acids throughout the CNS and periphery

Figure 6 illustrates a key finding in the study showing that Acute morphine significantly decreased levels of bile acids (CA, DCA, TCA) in the CNS and plasma without altering levels in liver and feces. Chronic treatment significantly increased levels of bile acids (DCA, TCA) throughout the CNS, plasma, liver, and feces. TCA (Figure 6A) signaling is largely unstudied in the CNS, so chromatograms of TCA standards (Figure 6B) and representative chromatograms are presented in Figures 6C (plasma) and 6D (cortex) of a 10 µL injection from partially purified 75% methanolic extracts of samples from mice that received chronic saline (grey) or chronic morphine (blue). Bar graphs in Figure 6E illustrate that that TCA levels were acutely lowered throughout the CNS, as well as the plasma. DCA’s chemical structure is shown in Figure 6F. As DCA signaling is similarly understudied in the CNS, we provide and chromatograms of DCA standards are displayed in Figure 6G, alongside chromatograms from a 10 uL injection of partially purified 100% methanolic extracts from of samples from mice that received chronic saline (grey) or chronic morphine (blue). As shown in representative chromatograms in Figures 6H and 6I, DCA was detected in plasma (6H) as well as the CTX (6I). Bar graphs in Figure 6J show that chronic morphine significantly increased DCA levels throughout the CNS, as well as in the plasma and liver. A heatmap summary of changes (Fig 7A) shows that where changes occurred in bile acid levels, acute morphine uniformly decreased levels. Conversely, chronic morphine uniformly increased levels where changes occurred. This is especially exemplified in the plasma where acute morphine decreased and chronic morphine increased levels of bile acids more than 3-fold.

## Discussion

In the present study we showed that Acute and Chronic morphine treatment causes distinct changes in a targeted set of endolipid signaling molecules. Many of these endolipids have not been fully characterized for their specific molecular targets (*i.e.* at receptors, channels, enzymes) and others, like the bile acids, are largely unstudied in the CNS, especially in the context of addiction. Data here provide information on novel endolipid signaling pathways that may contribute to the unwanted side effects of opioids, such as dependence and withdrawal. To illustrate how these data have the potential for discovery of novel signaling systems involved in opioid use, the following discussion focuses on the findings from four key classes of endolipids in the targeted screen; *N*-acyl GABAs (NAGABAs), *N*-acyl valines (NAVALs), *N*-acyl taurines (NATs), and specific bile acids.

### GABA signaling and opioids: how to the novel findings with N-acyl GABA endolipids play a role in Acute morphine signaling

Acute morphine administration significantly decreased levels of NAGABAs across the CNS. *N*-arachidonoyl GABA has antinociceptive properties when administered to rodents, though the mechanism of action for this behavioral finding is unknown [26, 27]. To the best of our knowledge, nothing is known about the link between NAGABAs and opioid signaling. However, activation of mu opioid receptors (MORs) is known to inhibit GABAergic neurotransmission in circuits that mediate analgesia and reward, though effects are region/circuit-specific and can diminish with chronic treatment [28, 29]. Therefore, other than the inclusion of GABA in the structure of NAGABAs, the relationship shown here between the two classes of compounds is novel. As an example, it is unknown if NAGABAs interact with GABA(A) or GABA(B) receptors. It is also unknown if NAGABAs act as a storage form for GABAs; however, depending on the brain region, GABA in the mouse brain is in µmole/g concentration range [30], whereas, here we show that CNS NAGABAs are measured in the picomoles/g range. Given the large discrepancy between concentrations, it is unlikely that NAGABAs contribute significantly to the GABA pool. The reverse may be more likely in that biosynthesis and regulation of NAGABAs are likely more sensitive to the levels of GABA available as a substrate, which in turn regulates NAGABA signaling, though testing this hypothesis is beyond the scope of this initial exploratory study. To date, there is still no clear understanding of the biosynthesis and metabolism of NAGABAs, with the exception that they are significantly decreased in the FAAH KO mouse brain [18], suggesting that FAAH is involved in their biosynthesis.

That NAGABAs were almost exclusively modulated in the Acute morphine paradigm here and not in the Chronic paradigm, suggests either an adaptation in this effect, or that it is only a transient effect and not observed 24 hours after exposure. In that NAGABAs are decreased with Acute morphine exposure, determining the pathways for NAGABA reduction in the CNS and this effect on neurophysiology could be a key in understanding additional ways in which morphine changes cellular signaling that is related to the pain reduction and euphoric phenotype that is associated with Acute opioid administration. Previous studies showed that members of the NAGABA family exhibit both agonist and antagonist activity at TRPV1 [17]. Additionally, NAGABA and its hydroxylated metabolite NAGABA-OH inhibit low-voltage gated T-type calcium channels, ion channels implicated in sleep and pain transmission [31, 32]. Understanding adaptation in signaling of NAGABAs at different targets that regulate neuroinflammation and neurotransmission (such as T-type calcium channels or TRPV channels) may be important for uncovering new non-opioid pathways engaged during initial drug treatment for therapeutic purposes that become dysregulated by chronic opioid use.

### N-acyl valines and opioids: novel link with chronic opioid exposure

Another prominent finding in the present study was that *N*-acyl valines (NAVALs) as a family were predominantly decreased by chronic opioid treatment throughout the CNS and periphery. Several NAVALs including *N*-stearoyl valine, *N*-oleoyl valine, *N*-linoleoyl valine, and *N*-docosahexaenoyl valine were previously characterized as antagonists of TRPV3 [17], a receptor enriched in keratinocytes whose overactivity can produce severe itching [33]. Interestingly, opioid analgesics can induce pruritis (excessive itching) in certain populations of patients [34]. Though other TRP channels (such as TRPV1) are known to regulate the activity of MORs (and vice versa), a direct relationship between chronic opioid and TRPV3 is not established [35, 36]. Given their antagonist effects on TRPV3, it is possible that lowering levels of NAVALs could increase TRPV3 activity and contribute to sensory/nociceptive dysregulation after chronic opioid use. Future studies that directly supplement NAVALs during withdrawal could determine if these behaviors could be modified.

Beyond its role in protein biosynthesis, valine as an amino acid potentially protects against oxidative stress and mitochondrial dysfunction [37]. This is relevant to the present study as chronic opioid administration results in oxidative stress and mitochondrial dysfunction in the CNS [38]. In this study, we observed that CNS levels of NAVALs were similar to plasma concentrations, which were several orders of magnitude lower than fecal levels. While the concentration gradient (feces>plasma>CNS) suggests that they could be acting either as nutrients or as signaling molecules influenced by gut microbes, fully characterizing this hypothesis is beyond the scope of the present study. Unpublished data from our lab shows that NAVALs are present and abundant in mouse chow. This finding suggests that the reduction of NAVALs in both the periphery and the CNS may be a result of the loss of appetite and increased intestinal transit associated with chronic opioid use in these mice, which may also serve as a useful biomarker for opioid withdrawal.

### Taurine signaling and opioids: new connections with N-acyl taurine endolipids

Taurine, a semi-essential amino acid that contains sulfur, is highly abundant in the central nervous system and plays a role in many physiological processes [39]. Supplementation of taurine has been investigated in preclinical settings as a strategy to reduce neuroinflammation associated with several neuropathological conditions including traumatic brain injury, ischemia, and neurodegenerative diseases [40-43]. More recently, the use of magnetic resonance spectroscopy (MRS) showed that chronic treatment with opioids lowers levels of taurine in the CNS in both rodents and non-human primates [44, 45]. Separate studies have observed that perinatal exposure to opioids causes lasting decreases in CNS taurine levels that persist through adolescence [46, 47]. Interestingly, treating opioid-dependent non-human primates with anti-withdrawal medications (methadone and clonidine) increased CNS taurine levels that were lowered in a state of opioid dependence [45]. Other than one study published more than 40 years ago showing that taurine attenuated naloxone-precipitated opioid withdrawal in mice [48], the physiological and behavioral characterization of taurine supplementation on unwanted side effects of opioids is largely unknown.

Here, we showed that chronic morphine consistently lowers levels of *N*-acyl taurine (NAT) levels across the CNS, with a preference for saturated (*N*-palmitoyl taurine and *N*-stearoyl taurine) and monounsaturated (N-oleoyl taurine) over the polyunsaturated *N*-arachidonoyl taurine. NATs are a family of endolipids metabolically regulated by the enzyme Fatty Acid Amide Hydrolase (FAAH), and signal at ion channels and G-Protein Coupled Receptors (GPCRs), including Transient Receptor Potential (TRP) channels, GPR119, potassium channels, and T-type calcium channels [32, 49-51]. NATs regulate diverse physiological processes, including inflammatory response via TRP channels, wound healing, and glucose homeostasis [49, 51, 52]. Among studies investigating the molecules included in the NAT family, *N*-arachidonoyl taurine has received the most attention. However, little is known about the signaling properties of the saturated NATs that were decreased by morphine throughout the CNS, including *N*-palmitoyl taurine and *N*-stearoyl taurine. To determine whether NATs play a role in symptoms of opioid withdrawal, a logical follow up to this observation would be to evaluate whether supplementation of NATs during chronic dosing reduces behavioral symptoms of spontaneous withdrawal.

### Bidirectional modulation of peripheral and CNS bile acids by acute and chronic morphine

In contrast to NATs that were decreased after chronic morphine, taurocholic acid (TCA), a taurine-conjugate of cholic acid, was markedly increased throughout the CNS and periphery. As a primary bile acid, cholic acid is synthesized in the liver via a pathway involving Cholesterol 7-alpha-hydroxylase (CYP7A1), and is conjugated to glycine or taurine through hepatic enzymes Bile Acid-CoA Synthetase (BACS) and Bile Acid-CoA:Amino Acid N-acyltransferase (BAAT) [53]. Of interest in the context of our study is a recently identified metabolic link between NATs and taurine-conjugated bile acids, showing that BAAT serves as a shared hepatic biosynthetic enzyme for both families [54]. The extent to which this occurs in the brain is unknown. Our working hypothesis is that the simultaneous lowering of NATs and increasing of TCA suggests that chronic morphine may shift metabolic processes regulating conjugation (biosynthesis) or metabolism of taurine-related molecules to favor some endolipid families (bile acids) over others (NATs). The way in which morphine affects the metabolic relationship between NATs and bile acids (especially in the CNS) provides a novel insight into the adaptations of morphine signaling and will benefit from future investigations.

TCA signals at GPCRs such as the Takeda G protein-coupled receptor 5 (TGR5, also known in literature as G-protein-coupled bile acid receptor or GPBAR1 or GPR19), and Sphingosine-1-Phosphate Receptor 2 (S1PR2) and to a lesser extent the nuclear Farsenoid X Receptor (FXR) [55]. TCA can act on TGR5 peripherally to stimulate GLP-1 release via TGR5 [56], a notable phenomenon given the prominent weight loss we saw in mice treated with chronic morphine. Recent lines of investigation suggest that TCA can act on receptors in the CNS. For example, TCA can lower blood pressure via TGR5 in the hypothalamus, while increased TCA promoted brain inflammation via S1PR2 in a rodent model of hepatic encephalopathy [57-59]. The role that TCA signaling via these receptors plays in neuroinflammatory dysregulation from chronic opioid exposure is largely unknown and the relationship between TCA regulation and morphine treatment shown here provides a clear validation for future studies.

In addition to TCA, Chronic morphine also increased the bile acid deoxycholic acid (DCA); however, it was also significantly decreased with Acute morphine and was the only endolipid in these screens to make the top 10 list of endolipids changed in both treatment paradigms. Deoxycholic acid is a metabolite of cholic acid and is one of many secondary bile acids that are produced by microbes in the gut [60]. As a signaling molecule, DCA was first identified in 1999 as an activator of FXR but has been recently shown to regulate neuroinflammation and mesolimbic neuronal activity in the CNS via TGR5 [61-63]. Additionally, *in vitro* studies demonstrate that DCA facilitates the enzymatic production of *N*-acylethanolamines (NAEs) by acting as an allosteric regulator of the enzyme *N*-acylphosphatidylethanolamine phospholipase-D (NAPE-PLD) thought the extent to which this occurs in vivo is unknown [64]. Morphine’s dynamic effects on peripheral DCA levels speak to the context-dependent relationship between opioids and DCA described in recent studies. For example, one group using subcutaneous morphine pellets (which deliver morphine steadily without peaks and troughs of injected levels) found that morphine decreased fecal DCA levels after 3 days of pellet implantation, an effect reversed by the MOR antagonist naltrexone [65]. In contrast, another study found increases in intestinal DCA after repeated intraperitoneal injections of fentanyl [66]. Given that gut dysbiosis and concurrent remodeling of the gut microbiome after chronic opioid exposure has been shown in recent years to play an important role in opioid reinforcement, bile acids may mediate some of these effects [67, 68].

To our knowledge, this is the first study to demonstrate that elevations in peripheral bile acids (such as TCA and DCA) after chronic opioid treatment correspond with bile acid elevation in the brain. Ursodeoxycholic acid (UDCA), a hydrophilic bile acid, is an FDA-approved treatment for cholestasis and can lower bile acid levels. UDCA and its taurine conjugated form, tauroursodeoxycholic acid (TUDCA), have been investigated as potential treatment strategies for neurodegenerative diseases such as Alzheimer’s [69, 70]. Promising medications exist to lower bile acids, so future studies that evaluate how the reduction of bile acids may be an additional tool that could be useful for treating certain aspects of opioid withdrawal if bile acid dysregulation directly contributes to opioid withdrawal.

## Conclusions and Future Directions

In the present study, using a targeted lipidomics approach, we identified several families of understudied endolipids that are dysregulated by opioid exposure, including NAGABAs, NAVALs, NATs, and bile acids. Given that morphine tended to decrease levels of endolipids in the brain, a natural follow up study could evaluate whether supplementation of endolipids (such as *N*-palmitoyl taurine or *N*-oleoyl valine) would ameliorate unwanted symptoms of chronic opioid exposure, such as withdrawal or analgesic tolerance. One challenge that arises is that our understanding of these endolipids’ signaling pathways is still incomplete. For example, while *N*-arachidonoyl taurine has been characterized as an activator of TRP channels, and inhibitor of T-type calcium channels, evidence about *N*-palmitoyl taurine’s effects on receptors/ion channels is largely unknown. Careful matching of these endogenous ligands to receptors/targets in the brain and body would provide a more complete understanding of non-opioid signaling pathways that may be useful in developing novel strategies for OUD.

## Acknowledgements

This project was supported by NIH grant P30DA056410 to HBB. Figure schematics created with Biorender.

## Author Contributions

TW and HBB designed the experiments. TW collected the data. TW, EOC, EFC, and CPS analyzed the data. TW and HBB wrote the manuscript with input from all authors. HBB and KM provided resources and supervised the project.

---

**Supplemental Figure 1:**
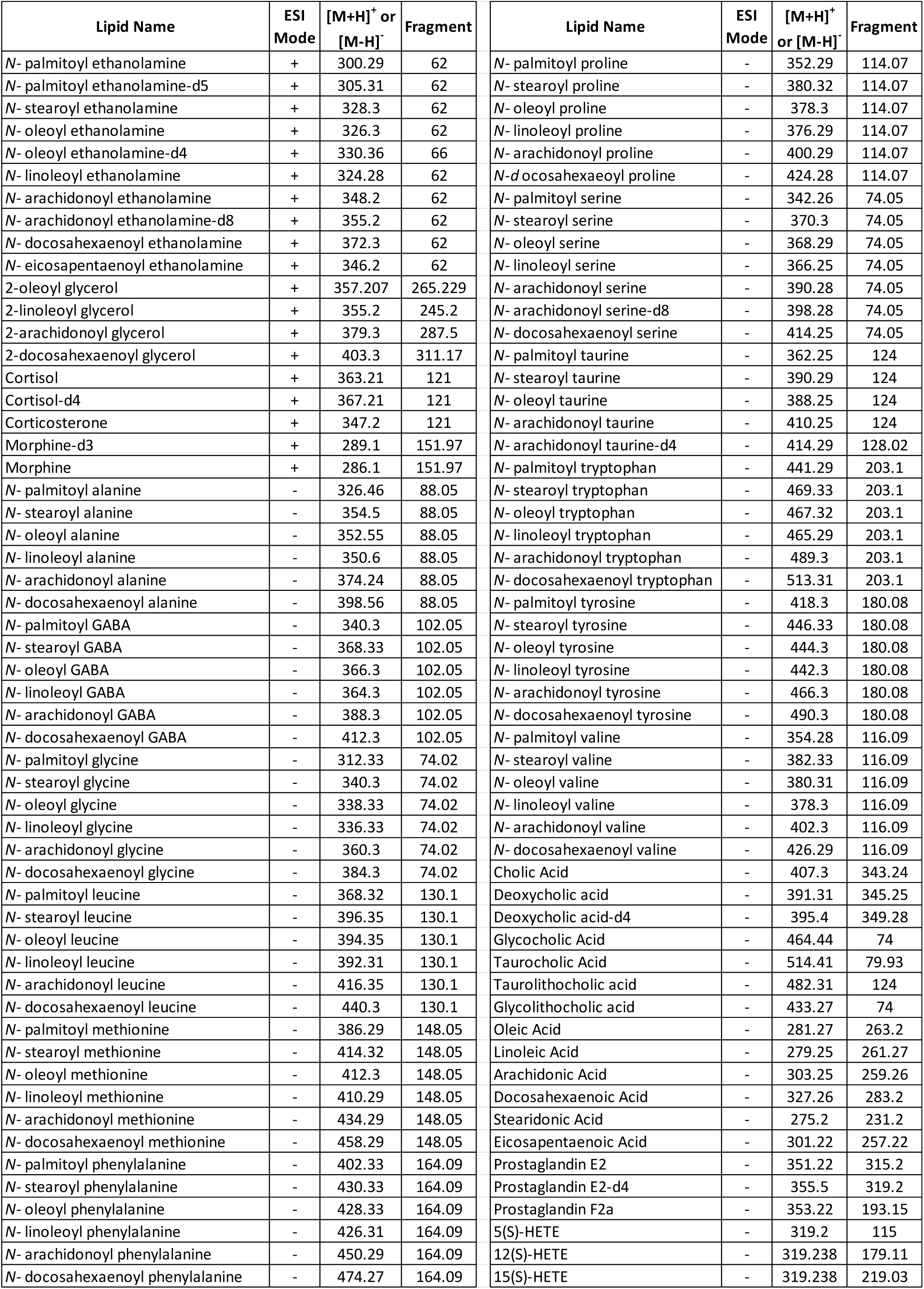
Full list of lipids in HPLC-MS/MS screening library. Lipids were screened from methanolic tissue extracts using a polarity switching multiple reactions monitoring (MRM) method. Lipid names are given with corresponding electrospray ionization (ESI) mode as well as parent/fragment m/z ratios.

**Supplemental Figure 2:**
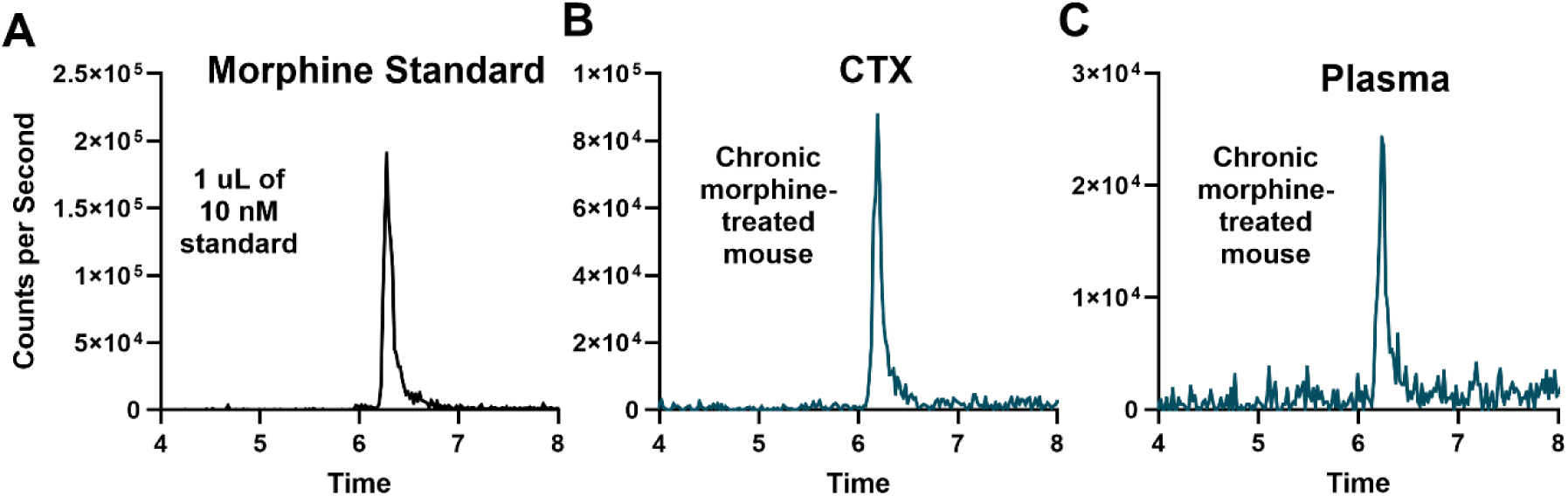
Detection of morphine in tissues 24 hours after final injection of chronic morphine. A) Representative chromatogram of a morphine standard (1 µL of a 10 nM solution) B) Representative chromatogram of morphine from partially purified methanolic extracts of the cortex (CTX) from a mouse treated with chronic morphine, dosed 24 hours prior to sacrifice C) Representative chromatogram of morphine in plasma from the same mouse.

**Supplemental Figure 3:**
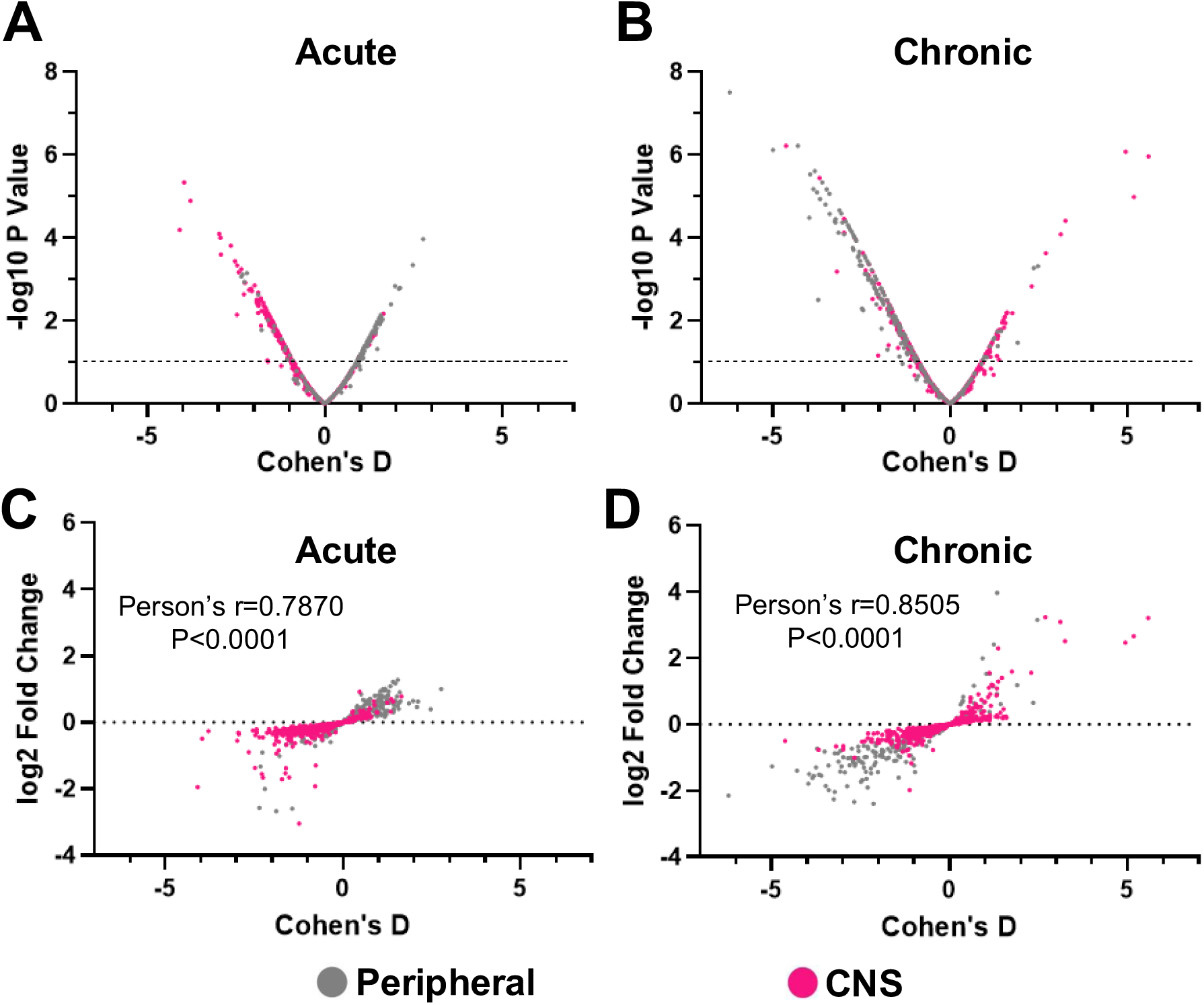
Relationships between Cohen’s D, p values and fold change. A) A plot displaying results from comparisons of Acute morphine treatment, with Cohen’s D values on the x axis and their corresponding –log10 P value for CNS regions (pink) and peripheral tissues (grey) on the y axis. B) A similar plot for comparisons of Chronic morphine treatment. C) A plot of Acute morphine comparisons displaying the correlation between Cohen’s D values on the x axis and log2 fold change on the y axis. D) A similar plot for comparisons of Chronic morphine.

**Supplemental Figure 4:**
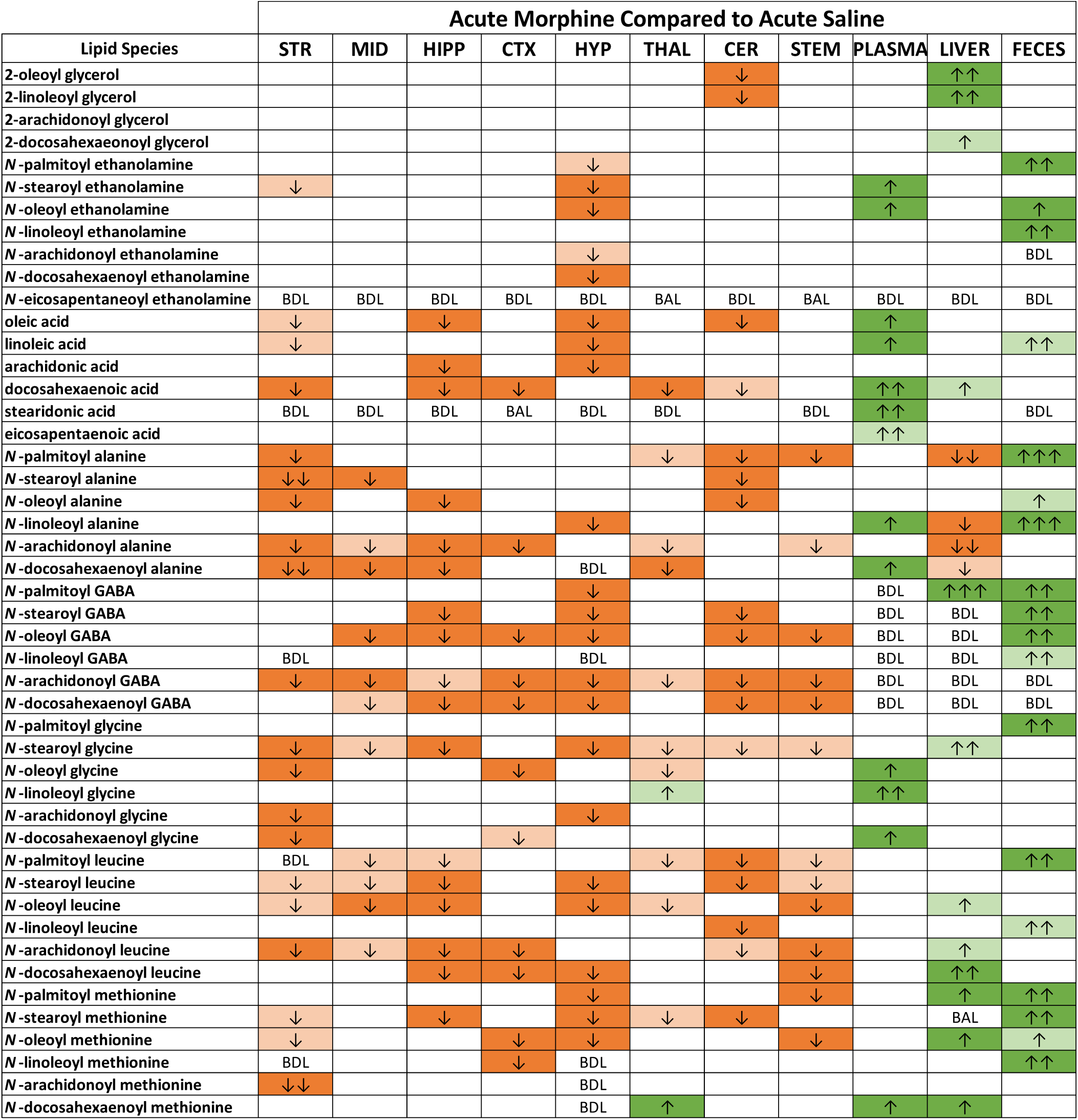
Acute morphine’s effects on lipid levels in the CNS and periphery. A summary heatmap displaying Acute morphine’s effect on the targeted lipidome in 8 brain regions in addition to plasma, liver, and feces. Heatmap key: Heatmaps were generated to summarize the effect of morphine on each lipid species in each tissue. Light green or light orange boxes depict a P value between 0.1 and 0.05, while dark green or dark orange depict a P value below 0.05. Arrows summarizing fold change of morphine compared to saline are presented as an increase(↑) or decrease(↓). Number of arrows correspond with magnitude of fold change. One arrow: 1-1.49 fold difference; two arrows: 1.5-1.99 fold difference; three arrows 2-2.99 fold difference; four arrows 3-9.99 fold difference, five arrows 10+ fold difference.

**Supplemental Figure 5:**
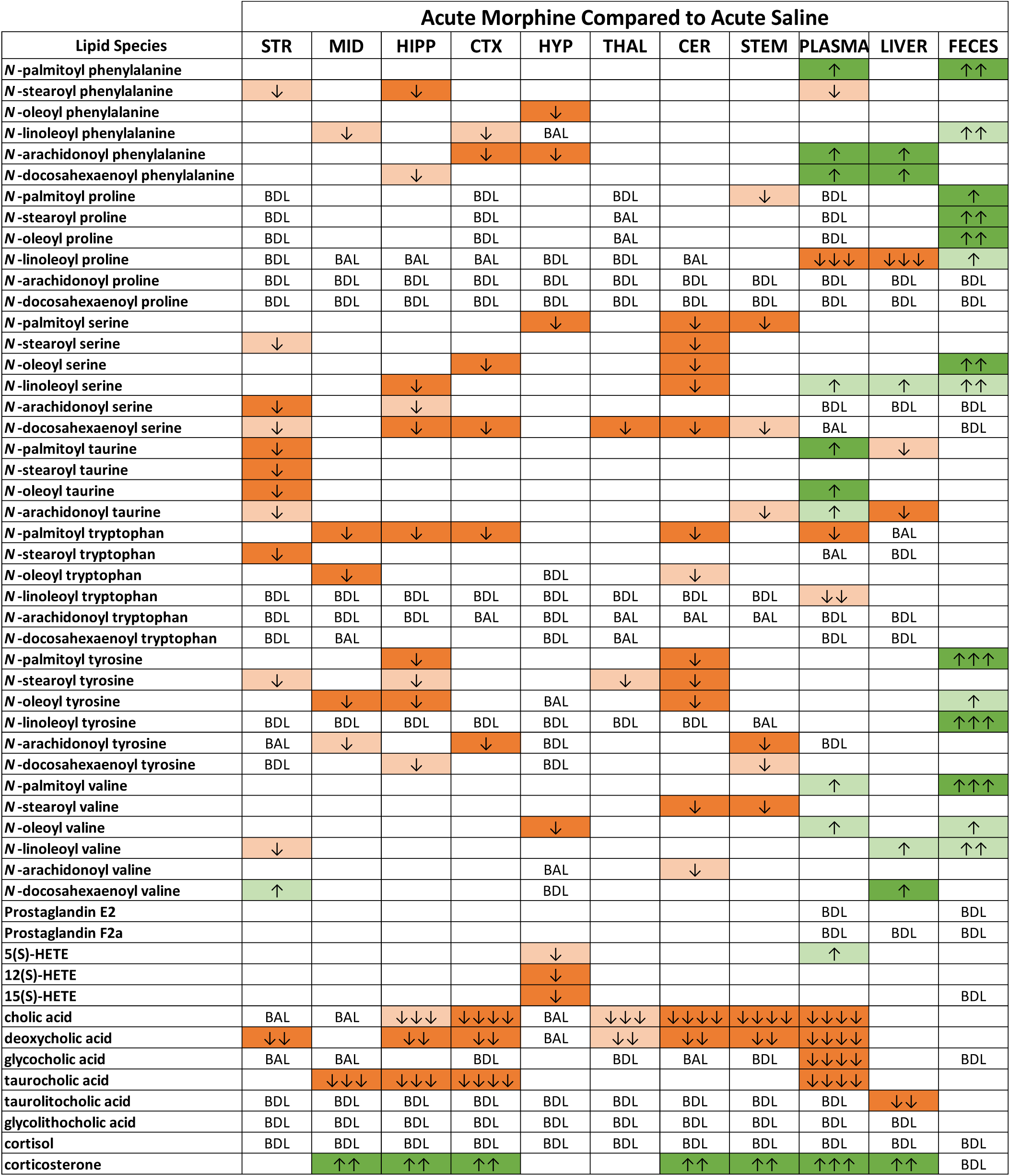
Acute morphine’s effects on lipid levels in the CNS and periphery. A summary heatmap displaying Acute morphine’s effect on the targeted lipidome in 8 brain regions in addition to plasma, liver, and feces.

**Supplemental Figure 6:**
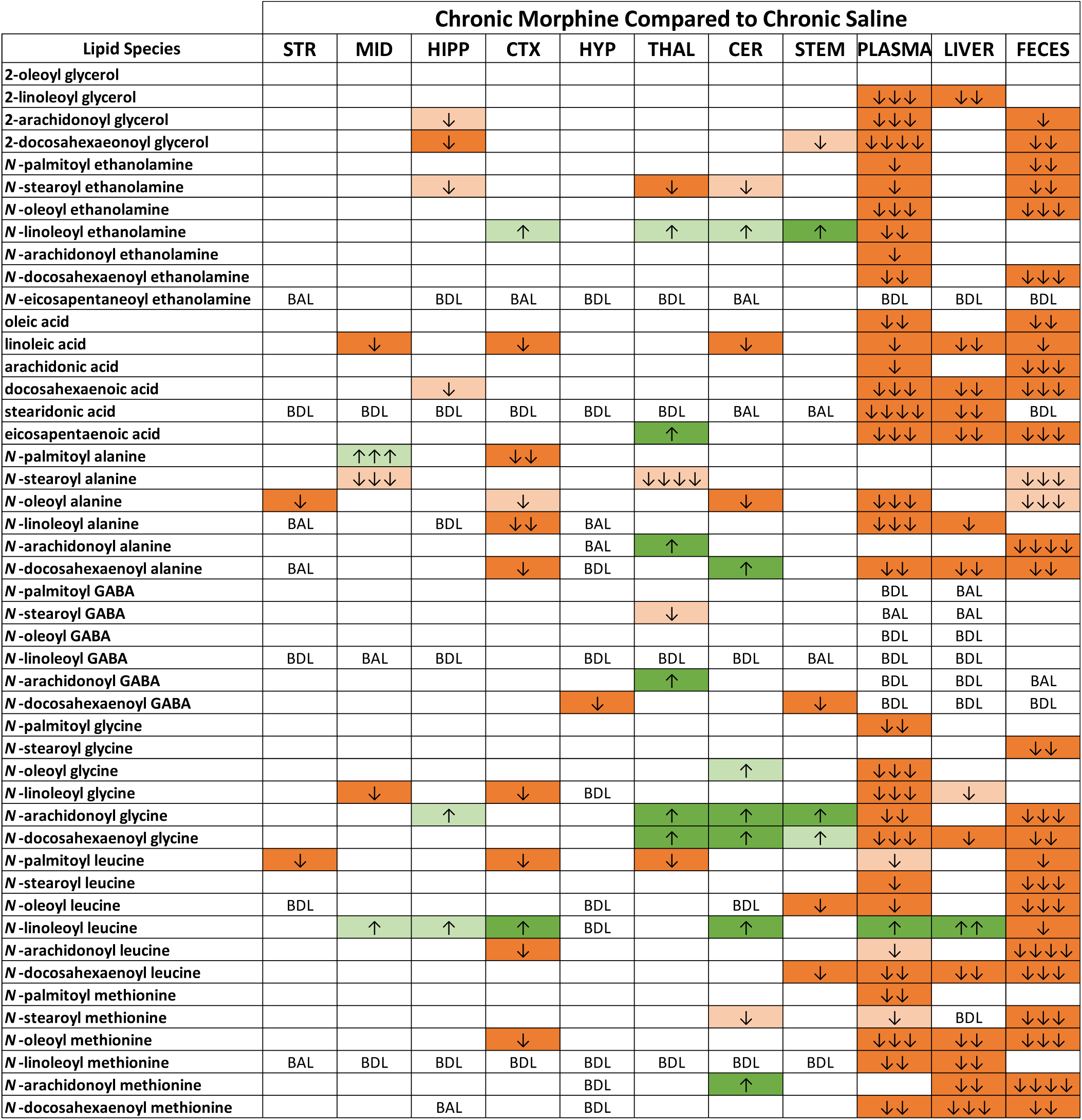
Chronic morphine’s effects on lipid levels in the CNS and periphery. A summary heatmap displaying effect of Chronic morphine on the targeted lipidome in 8 brain regions in addition to plasma, liver, and feces.

**Supplemental Figure 7:**
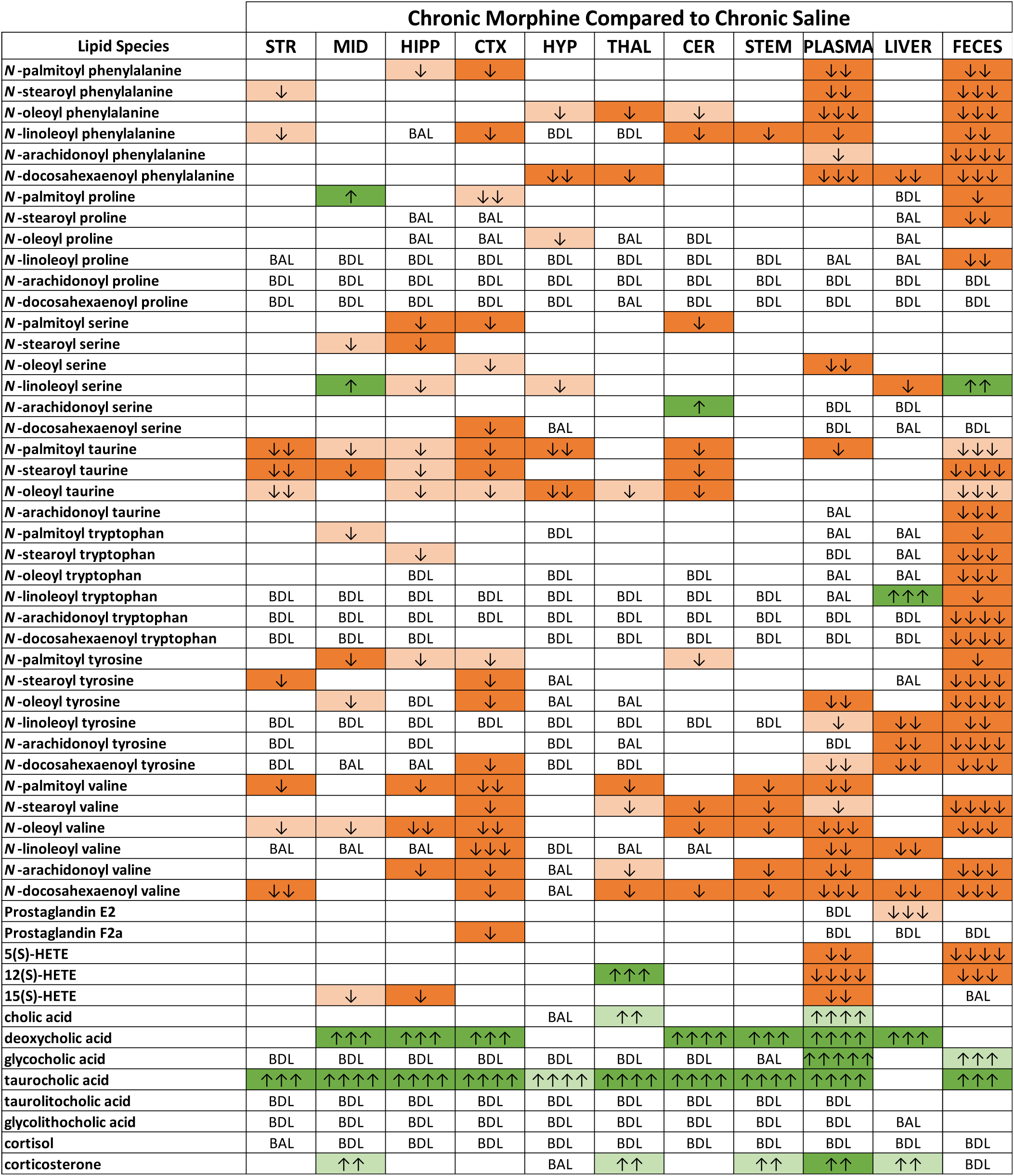
Chronic morphine’s effects on lipid levels in the CNS and periphery. A summary heatmap displaying effect of Chronic morphine on the targeted lipidome in 8 brain regions in addition to plasma, liver, and feces.

**Supplemental Figure 8:**
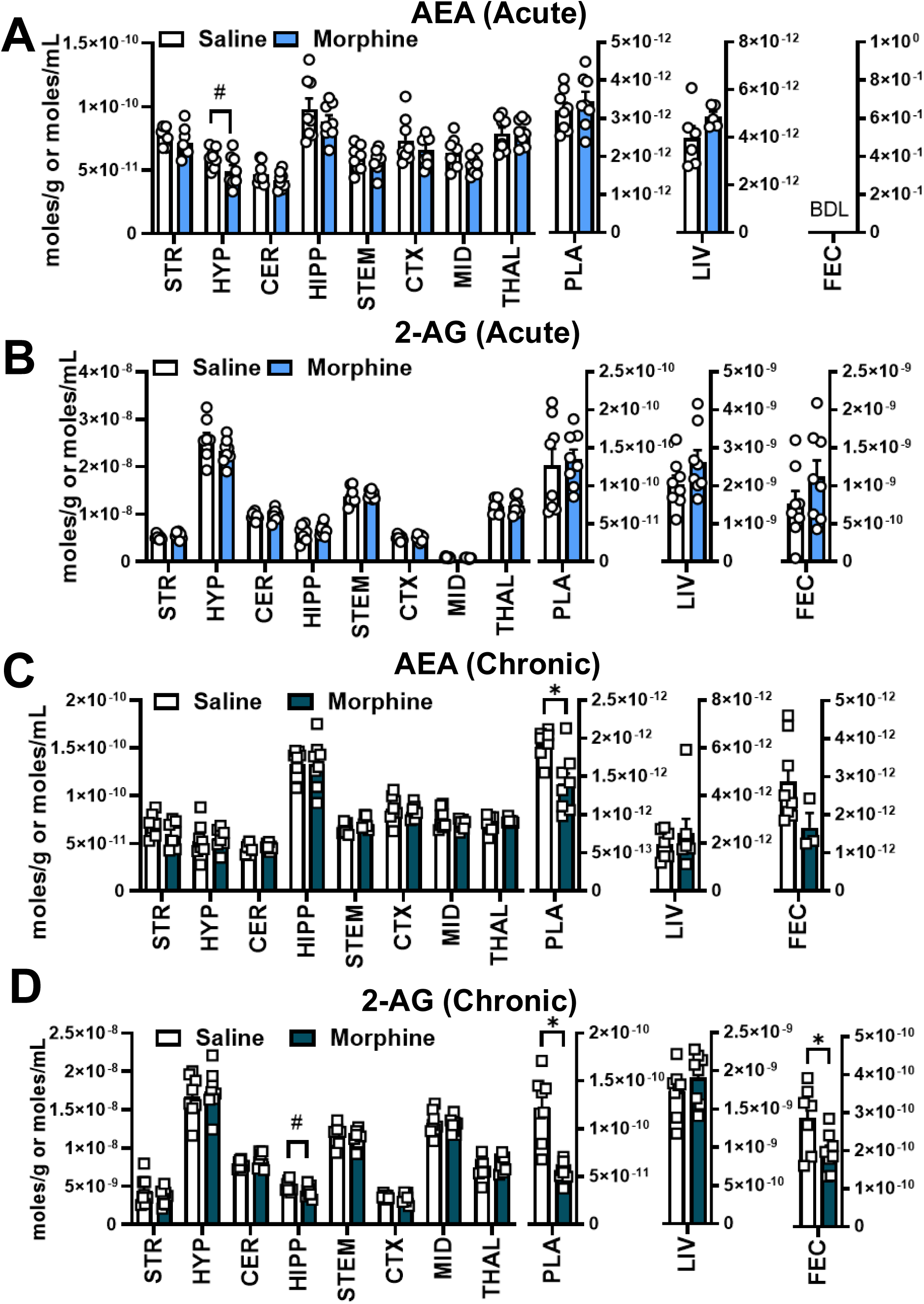
Effect of Acute and Chronic morphine administration on levels of endocannabinoids. A) Levels of Anandamide (AEA) in the CNS and peripheral tissues after acute administration of saline or morphine. B) Levels of 2-arachidonoyl glycerol (2-AG) in the CNS and peripheral tissues after acute administration of saline or morphine. C) Levels of Anandamide (AEA) in the CNS and peripheral tissues after chronic administration of saline or morphine. D) Levels of 2-arachidonoyl glycerol (2-AG) in the CNS and peripheral tissues after chronic administration of saline or morphine. K) Two tailed t-test: #=p value between .05 and .01, *=p<.05. Data are presents as the mean +/-SEM with n=8 per treatment group. Tissue Abbreviations: STR=striatum, MID=midbrain, CTX=cortex, HYP= hypothalamus, STEM=brainstem, THAL=thalamus, CER= cerebellum, PLA=plasma, LIV=liver, FEC=feces

